# Mosquito-bacteria interactions during larval development trigger metabolic changes with carry-over effects on adult fitness

**DOI:** 10.1101/2021.05.20.444942

**Authors:** Émilie Giraud, Hugo Varet, Rachel Legendre, Odile Sismeiro, Fabien Aubry, Stéphanie Dabo, Laura B. Dickson, Claire Valiente Moro, Louis Lambrechts

## Abstract

In animals with distinct life stages such as holometabolous insects, adult phenotypic variation is often shaped by the environment of immature stages, including their interactions with microbes colonizing larval habitats. Such carry-over effects were previously observed for several adult traits of the mosquito *Aedes aegypti* after larval exposure to different bacteria, but the mechanistic underpinnings are unknown. Here, we investigated the molecular changes triggered by gnotobiotic larval exposure to different bacteria in *Ae. aegypti*. We initially screened a panel of 16 bacterial isolates from natural mosquito breeding sites to determine their ability to influence adult life-history traits. We subsequently focused on four bacterial isolates (belonging to *Flavobacterium, Lysobacter, Paenibacillus*, and *Enterobacteriaceae*) with significant carry-over effects on adult survival and found that they were associated with distinct transcriptomic profiles throughout mosquito development. Moreover, we detected carry-over effects at the level of gene expression for the *Flavobacterium* and *Paenibacillus* isolates. The most prominent transcriptomic changes in gnotobiotic larvae reflected a profound remodeling of lipid metabolism, which translated into phenotypic differences in lipid storage and starvation resistance at the adult stage. Together, our findings indicate that larval exposure to environmental bacteria trigger substantial physiological changes that impact adult fitness, uncovering a mechanism underlying carry-over effects of mosquito-bacteria interactions during larval development.

## Introduction

The life cycle of holometabolous insects is characterized by complete metamorphosis of the final immature stage into a markedly distinct mature stage. The holometabolous life cycle may have evolved to prevent larvae from competing with adults because they generally inhabit different ecological niches [1]. The environment of immature stages, however, has a profound influence on adult life history through various ‘carry-over effects’ [2, 3]. Both abiotic factors (e.g., temperature, nutrient availability) and biotic interactions (e.g., predation, competition, mutualism) of the larval environment contribute to determine adult traits [4–7].

Mosquitoes are holometabolous insects of particular importance for public health because they serve as vectors for several human pathogens. For instance, *Aedes aegypti* is a widely distributed mosquito species [8] whose recent evolution fueled the emergence of dengue virus (DENV), Zika virus (ZIKV) and other medically important arthropod-borne viruses [9, 10]. Studies in *Ae. aegypti* and other mosquitoes have shown that the larval environment can significantly influence adult life history and physiology. For example, larval rearing temperature, diet and crowding determine body size, longevity, energy reserves and immune function at the adult stage [11–15]. In addition, the larval environmental can affect the ability of adult mosquitoes to acquire and transmit arboviruses (i.e., vector competence) [16]. For instance, larval competition [17, 18], food availability [19], rearing temperature [20, 21], and exposure to insecticides [22] can modulate vector competence at the adult stage, although these effects are often complex and interdependent.

In recent years, it has become clear that environmental microbes colonizing mosquito larval habitats play a major role in both larval nutrition and development [23, 24]. Successful adult emergence critically depends on larval exposure to live bacteria [25–27], although supplementing axenic (i.e., germ-free) mosquito larvae with a specific diet can partially restore development [28]. The mechanisms underlying the effects of mosquito-microbiota interactions on immature development remain to be fully elucidated. It has been proposed that bacteria in the larval gut provide the essential micronutrient riboflavin (vitamin B2), whose lack results in gut hypoxia and developmental arrest [29–31]. Other studies suggested that the role of bacteria during larval development is primarily nutritional [28], possibly by contributing to folate biosynthesis and/or by enhancing energy storage [32]. This is consistent with work in *Drosophila* showing that larval gut bacteria cooperate to establish an integrated nutritional network supporting host growth [33, 34].

Mosquito-bacteria interactions at the larval stage can also impact adult traits via carry-over effects. In a previous study, we found that *Ae. aegypti* exposure to different natural bacterial isolates during larval development resulted in differences in adult body size, antibacterial activity and DENV vector competence [35]. Given that the bacterial microbiota of wild larvae varies substantially between natural microhabitats [25, 35–37], mosquitoes developing in different breeding sites are thus expected to contribute differentially to pathogen transmission. To date, the mechanism(s) underlying such carry-over effects of mosquito-bacteria interactions at the larval stage are unknown. Of note, we previously observed that larval exposure to bacteria mediated carry-over effects in the absence of trans-stadial transfer of the bacteria through metamorphosis [35]. This is consistent with another study in which larval exposure to bacteria modulated infection by DENV and ZIKV in adult *Ae. aegypti* in the absence of bacterial transfer between the life stages [38]. These observations support the hypothesis that carry-over effects result from indirect consequences of mosquito-bacteria interactions at the larval stage through physiological changes of the host that persist trans-stadially.

Here, we investigated the physiological changes associated with carry-over effects of larval exposure to bacteria in *Ae. aegypti*. We screened a collection of bacteria isolated during our previous study [35] for carry-over effects on adult phenotypes. Larvae were reared under gnotobiotic conditions in mono-association (i.e., in presence of a single bacterial isolate) until the pupal stage, after which adult mosquitoes were maintained and tested under standard (non-sterile) insectary conditions. We focused on four bacterial isolates that mediated significant carry-over effects on adult life-history traits and compared the transcriptomic profiles of larvae, pupae and adults between the four gnotobiotic treatments. Our results indicate that exposure to different bacteria during larval development is associated with major transcriptomic changes, some of which reflect a profound remodeling of lipid metabolism with functional repercussions on adult fitness.

## Results

### Larval exposure to different bacteria influences life-history traits

We previously isolated a panel of bacteria from the water of natural *Ae. aegypti* breeding sites in both domestic (D) and sylvatic (S) habitats in Gabon [35]. Comparing domestic and sylvatic isolates was not a primary goal of the present study and we denote isolates with D or S thereafter mainly for identification purposes. We first screened 16 of these bacterial isolates (representing the taxonomic diversity of bacterial genera across sites and habitats) for their ability to influence mosquito life-history traits through interactions during larval development. We compared the developmental time, adult female body size and adult female survival of mosquitoes after mono-association of gnotobiotic larvae with each one of the 16 bacterial isolates. We generated gnotobiotic larvae by exposing axenic larvae to a standardized concentration of a single bacterial isolate in otherwise sterile conditions, and used non-axenic larvae (i.e., from non- sterilized eggs) as controls. Developmental time was evaluated by monitoring the proportion of pupating individuals over time. The proportion of larvae that successfully pupated ranged from 70% to 100% across gnotobiotic treatments, experiments and replicates but pupation success was high overall (mean: 95.7%; median 100%). The median developmental time (50% pupation day; PD_50_) varied significantly (*p*<0.0001) between gnotobiotic treatments (Fig. 1A), ranging from 8.03 days (*Lysobacter* D) to 11.62 days (*Rahnella* S). The PD_50_ was 8.69 days for non-axenic controls. The body size of adult females also varied significantly (*p*<0.0001) between gnotobiotic treatments (Fig. 1B). The mean female wing length (normalized to account for experimental variation) ranged from –0.041 mm (*Leifsonia* S) to +0.254 mm (*Rahnella* S) relative to the non-axenic controls. Accounting for differences between experiments, the gnotobiotic treatment influenced adult female survival relative to the non-axenic controls for seven of the 16 bacterial isolates (Fig. 1C; Table S1). Together, these experiments showed that exposure to different bacteria during larval development resulted in variation in several life-history traits, including adult female survival, a critical determinant of vectorial capacity [39]. We selected the three isolates with the strongest effect on adult female survival (*Flavobacterium* S, *Lysobacter* D and *Paenibacillus* D) and a fourth isolate (*Enterobacteriaceae* S) that we previously found to influence adult antibacterial activity and DENV vector competence [35], for further study.

**Fig. 1.**
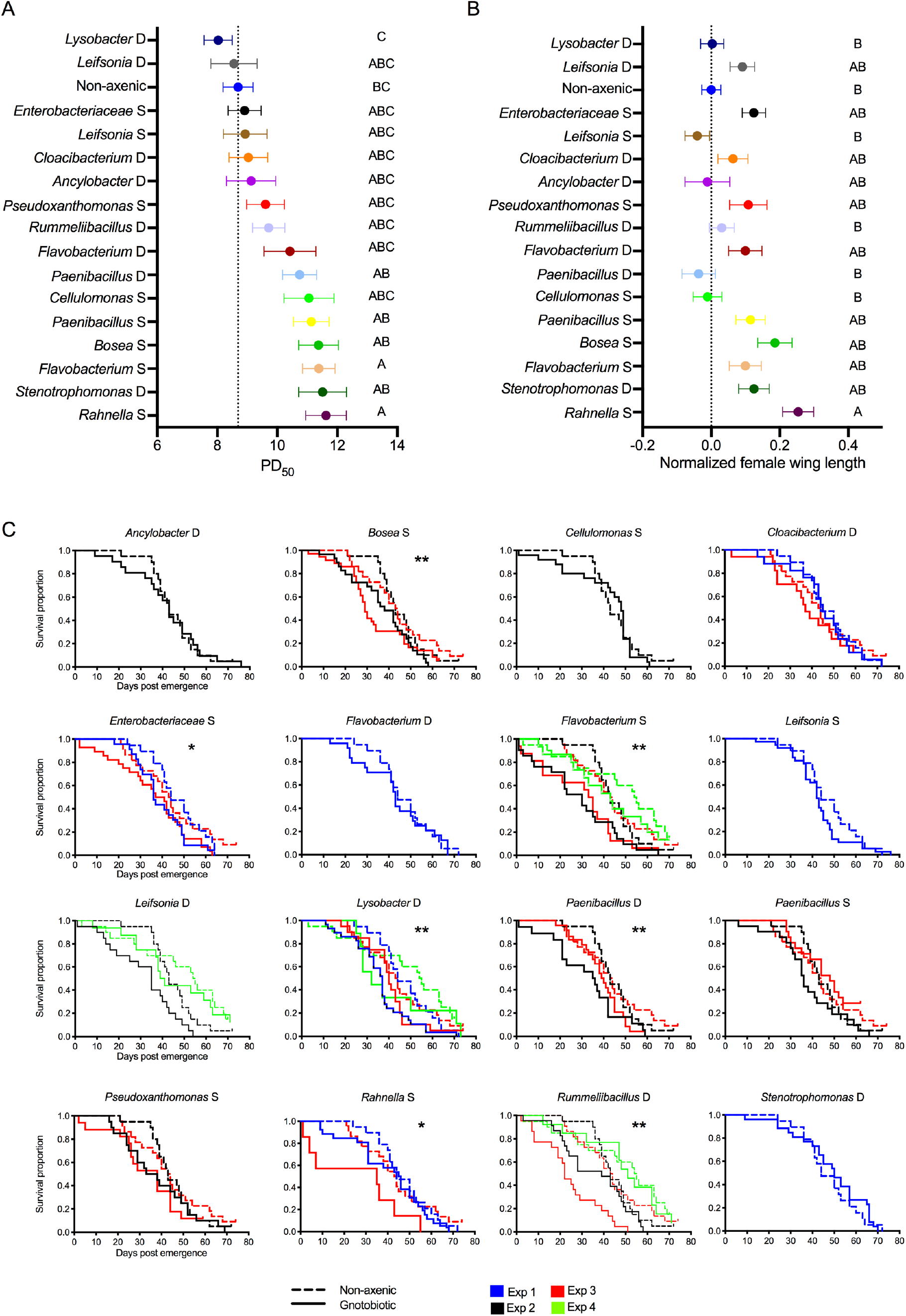
Larval exposure to different bacterial isolates results in variation in several life-history traits. The figure shows *Ae. aegypti* variation in developmental time (**A**), female wing length (**B**) and female adult survival (**C**) following larval exposure to one of 16 bacterial isolates. **A**. The forest plot shows the 50% pupation day (PD_50_) estimate (number of days elapsed until 50% of larvae have become pupae) for each of the 16 bacterial isolates, weighted by the sample size (number of pupating individuals per replicate). Separate PD_50_ estimates were obtained for each replicate flask with a Kaplan-Meier analysis of all pupating individuals. The PD_50_ estimates were compared between treatments by weighted ANOVA. **B**. The forest plot shows the normalized wing length of adult females for each of the 16 bacterial isolates. The wing length was normalized relative to the average wing length of the non-axenic controls from the same experiment. The normalized wing length was compared between treatments by one-way ANOVA. The experiment effect was removed from the model because it was non-significant (*p*=0.1022). In panels **A**-**B**, data represent the mean ± SEM from three independent experiments in triplicate. Bacterial isolates are ordered according to their PD_50_ estimate. The vertical dotted line represents the mean value of the non-axenic controls. Statistical significance of the pairwise differences was determined by Tukey’s post-hoc test and is indicated by letters next to the graphs. Treatments not connected by the same letter are significantly different. **C**. Kaplan-Meier plots showing adult female survival over time in four separate experiments for each of the 16 bacterial isolates. The dotted lines represent the non-axenic controls and the solid lines represent the gnotobiotic treatment. Asterisks indicate the statistical significance of the main treatment effect (\**p*<0.05, \*\**p*<0.01). The full statistical analyses of survival curves are provided in Table S1. In all panels the bacterial isolate is identified by the genus name followed by the letter D or S to denote isolation from a domestic or sylvatic larval habitat, respectively.

### Larval exposure to different bacteria influences DENV vector competence

We first examined whether the bacterial load differed between the four gnotobiotic treatments. We found that the amount of bacterial DNA (quantified by the normalized concentration of the universal *16S* bacterial ribosomal RNA gene) was similar in larvae across gnotobiotic treatments although it was significantly lower than in the non-axenic controls (Fig. 2A). The bacterial load did not significantly vary in pupae (Fig. 2B) or in adults shortly after emergence under sterile conditions (Fig. 2C). We also monitored the concentration of cultivable bacteria in the larval rearing water and found that it was similar between gnotobiotic treatments at early time points and significantly lower at late time points for *Lysobacter* D, *Paenibacillus* D, and *Enterobacteriaceae* S isolates (Fig. S1). After gnotobiotic rearing followed by 5-7 days spent under conventional insectary conditions, we offered adult *Ae. aegypti* females a DENV infectious blood meal. We found that the proportion of mosquitoes that became infected was significantly lower for females originating from larvae mono-associated with *Flavobacterium* S or *Paenibacillus* D compared to the non-axenic controls (Fig. 2D). The proportion of DENV-infected females that developed a disseminated infection did not differ between treatments (Fig. 2E), however the titer of disseminated virus was significantly lower (about 5-fold) in mosquitoes whose larvae were exposed to *Flavobacterium* S and *Paenibacillus* D (Fig. 2F). The titer of disseminated virus is a proxy for DENV transmission potential in *Ae. aegypti* [40]. Thus, these experiments showed that despite harboring a similar bacterial load, mosquitoes exposed to different bacteria during larval development had a different ability to transmit DENV at the adult stage.

**Fig. 2.**
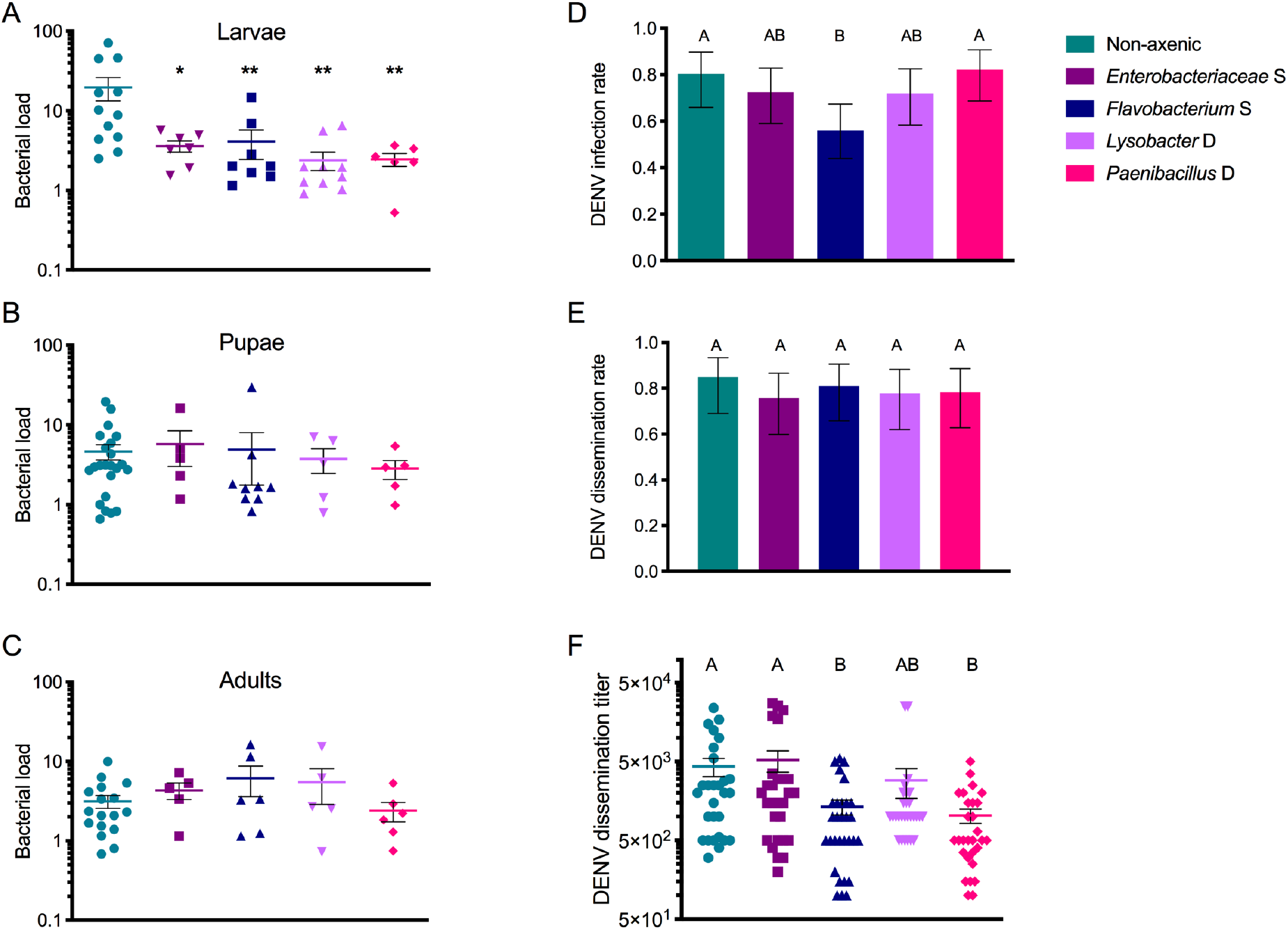
Larval exposure to four selected bacterial isolates results in variation in DENV vector competence. The figure shows bacterial loads in individual L_4_ larvae (**A**), pupae (**B**) and freshly emerged adults (**C**) and dengue virus (DENV) infection rate (**D**), dissemination rate (**E**) and dissemination titer (**F**) after larval exposure to *Enterobacteriaceae* S, *Flavobacterium* S, *Lysobacter* D or *Paenibacillus* D isolates. In panels **A**-**C**, the bacterial load was measured by quantitative PCR using primers targeting a conserved sequence in the *16S* bacterial rRNA gene and normalized to the mosquito housekeeping gene *RP49*. The horizontal bar indicates the mean and the error bar represents the SEM. Asterisks indicate statistically significant differences between non-axenic and gnotobiotic treatments (\**p*<0.05, \*\**p*<0.01) by Mann– Whitney *U* test. In panels **D**-**E**, error bars represent the 95% confidence intervals of the proportions. The infection rate is the proportion of blood-fed females with a DENV-positive body 14 days post infectious blood meal. The dissemination rate is the proportion of infected females with a DENV-positive head 14 days post infectious blood meal. In panel **F**, the dissemination titer is the concentration of infectious DENV particles expressed as the log_10_-transformed number of focus-forming units (FFU) detected in the head 14 days post infectious blood meal. The horizontal bar indicates the mean and the error bar represents the SEM. Panels **D**-**F** represent data from two separate experiments that were analyzed by nominal logistic regression (**D**-**E**) or by ANOVA (**F**) as a function of experiment, treatment and their interaction. In panel **D**, the treatment effect was statistically significant (*p*=0.0399) whereas the experiment effect (*p*=0.0597) and the interaction effect (*p*=0.9559) were not. In panel **E**, there was no statistically significant effect of the treatment (*p*=0.8984), the experiment (*p*=0.0849) or the interaction (*p*=0.9772). In panel **F**, the treatment effect (*p*<0.0001) and the experiment effect (*p*=0.0203) were statistically significant whereas the interaction effect (*p*=0.1798) was not. In panels **D**-**F**, the statistical significance of pairwise differences between treatments are indicated by letters above the graphs (based on the 95% confidence intervals in panels **D**-**E** and on Tukey’s post-hoc test in panel **F**). Treatments not connected by the same letter are significantly different.

### Larval exposure to different bacteria results in substantial transcriptional changes

We next investigated whether exposure to different bacteria at the larval stage was accompanied by differences in transcriptional profiles throughout development. We sequenced the whole-body transcriptomes of triplicate pools (N=12 per pool) of L_4_ larvae, pupae and newly emerged adults (6 males + 6 females) from gnotobiotic and non-axenic treatments. We also included pools of 12 L_1_ axenic larvae as a reference but our primary aim was to compare gnotobiotic treatments between them and with non-axenic controls. Across samples and life stages, we detected a total of 19,763 *Ae. aegypti* transcripts. Principal component analysis (PCA) of read counts identified three main clusters representing the three life stages (Fig. 3A). Of note, the larval cluster included a few pools of pupae, the pupal cluster contained a few pools of larvae and adults, and the adult cluster included a few pools of pupae. There was no overlap between pools of larvae and pools of adults. These results are consistent with the existence of a transcriptional program specific to each life stage, with a potential mismatch with the apparent morphology during transitioning periods. There was a general tendency for pools from the same experimental treatment to cluster together in the PCA. When larval samples were analyzed alone, the L_1_ axenic larvae clustered separately from the rest and the gnotobiotic larvae from the *Lysobacter* D and *Flavobacterium* S treatments did not overlap with the non-axenic controls (Fig. 3B). Pairwise comparisons found a total of 2,359, 10,796 and 6,026 transcripts that were significantly differentially expressed between at least two gnotobiotic treatments in larvae, pupae and newly emerged adults, respectively. To facilitate comparisons across the four gnotobiotic treatments, we used the non-axenic controls as a reference in further analyses. A total of 4,029, 7,240 and 2,844 transcripts were significantly up- or down-regulated relative to the non-axenic controls in gnotobiotic larvae, pupae and newly emerged adults, respectively. Only a small minority of differentially expressed genes was shared among the four gnotobiotic treatments, for each life stage (Fig. 3C). Functional clustering revealed that the most represented gene ontology (GO) category in differentially expressed genes was that of metabolic processes, irrespective of the life stage (Fig. 3D). Pathway enrichment analysis of up-regulated transcripts confirmed that metabolic pathway was the most represented Kyoto Encyclopedia of Genes and Genomes (KEGG) category in all of the comparisons (Fig. S2). Together, the transcriptomic analyses showed that exposure to different bacteria at the larval stage resulted in massive transcriptional changes, most of which were related to metabolic processes.

**Fig. 3.**
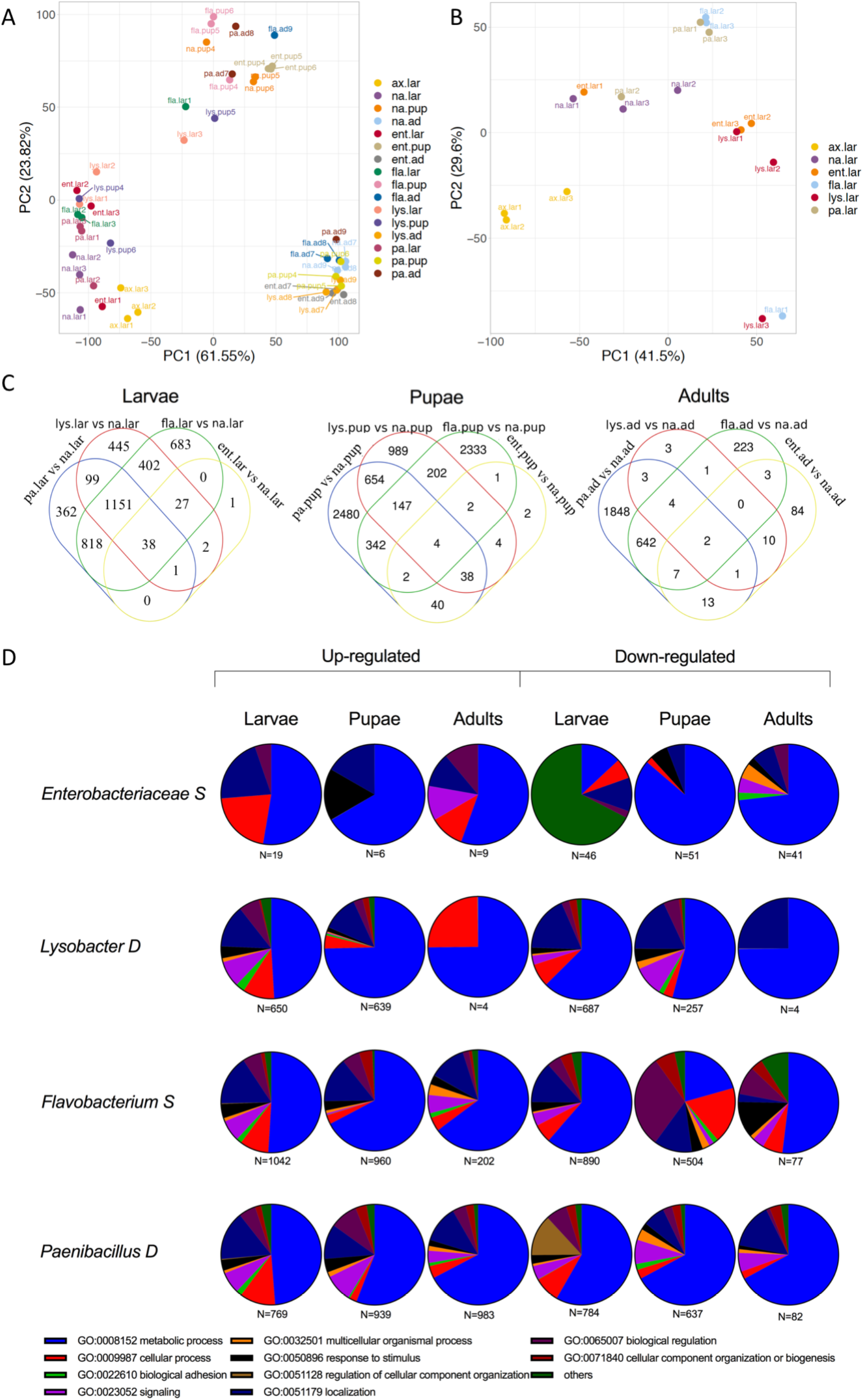
Larval gnotobiotic treatments trigger profound transcriptomic changes throughout mosquito development. The figure summarizes the transcriptional profiles of L_4_ larvae, pupae and newly emerged adults (triplicate pools of 12 individuals for each life stage, with 6 males + 6 females at the adult stage) after larval exposure to four selected bacterial isolates, or non-axenic control treatment. Triplicate pools of axenic L_1_ larvae are also included. **A**. Principal component analysis (PCA) of read counts among the 48 experimental groups (3 life stages 5 treatments 3 replicates + axenic larvae in triplicate). **B**. PCA of read counts in larvae only. In panels **A**-**B**, the PCA is based on the variance-stabilized transformed count matrix and only the first two components are shown. The percentage of variability explained by each component is displayed in brackets. The data points are identified by the combination of the isolate (ent=*Enterobacteriaceae* S; fla=*Flavobacterium* S; lys=*Lysobacter* D; pa=*Paenibacillus* D; na=non-axenic) and the life stage (lar=larvae; pup=pupae and ad=adults), followed by the replicate number. **C**. Venn diagrams showing the overlap in the number of differentially expressed transcripts (relative to the non-axenic controls) between the four gnotobiotic treatments in larvae, pupae and newly emerged adults. **D**. Functional clustering of transcripts differentially expressed relative to non-axenic controls, for each combination of life stage (in columns) and gnotobiotic treatment (in rows). Pie charts show the proportion of gene ontology (GO) categories for up- and down-regulated transcripts. GO categories with <1% of differentially expressed transcripts in all comparisons are grouped together in the category designated as ‘others’. N indicates the total number of transcripts assigned to functional categories.

### Carry-over effects occur at the level of transcriptional regulation

We next asked whether we could detect transcripts whose differential expression was correlated between life stages. For each gnotobiotic treatment, we defined ‘carry-over transcripts’ as transcripts that were (*i*) differentially regulated in larvae, (*ii*) differentially regulated in adults, and (*iii*) differentially regulated in the same direction and with a similar magnitude in both larvae and adults (i.e., a lack of statistical interaction). We identified 1, 180 and 128 such ‘carry-over transcripts’ in the *Lysobacter* D, *Flavobacterium* S and *Paenibacillus* D treatments, respectively (Fig. 4A; Fig. S3). Functional clustering revealed that the most represented GO category in ‘carry-over transcripts’ of the *Flavobacterium* S and *Paenibacillus* D treatments was that of metabolic processes (Fig. 4B). This analysis showed that carry-over effects observed at the phenotypic level can also be detected at the level of gene expression.

**Fig. 4.**
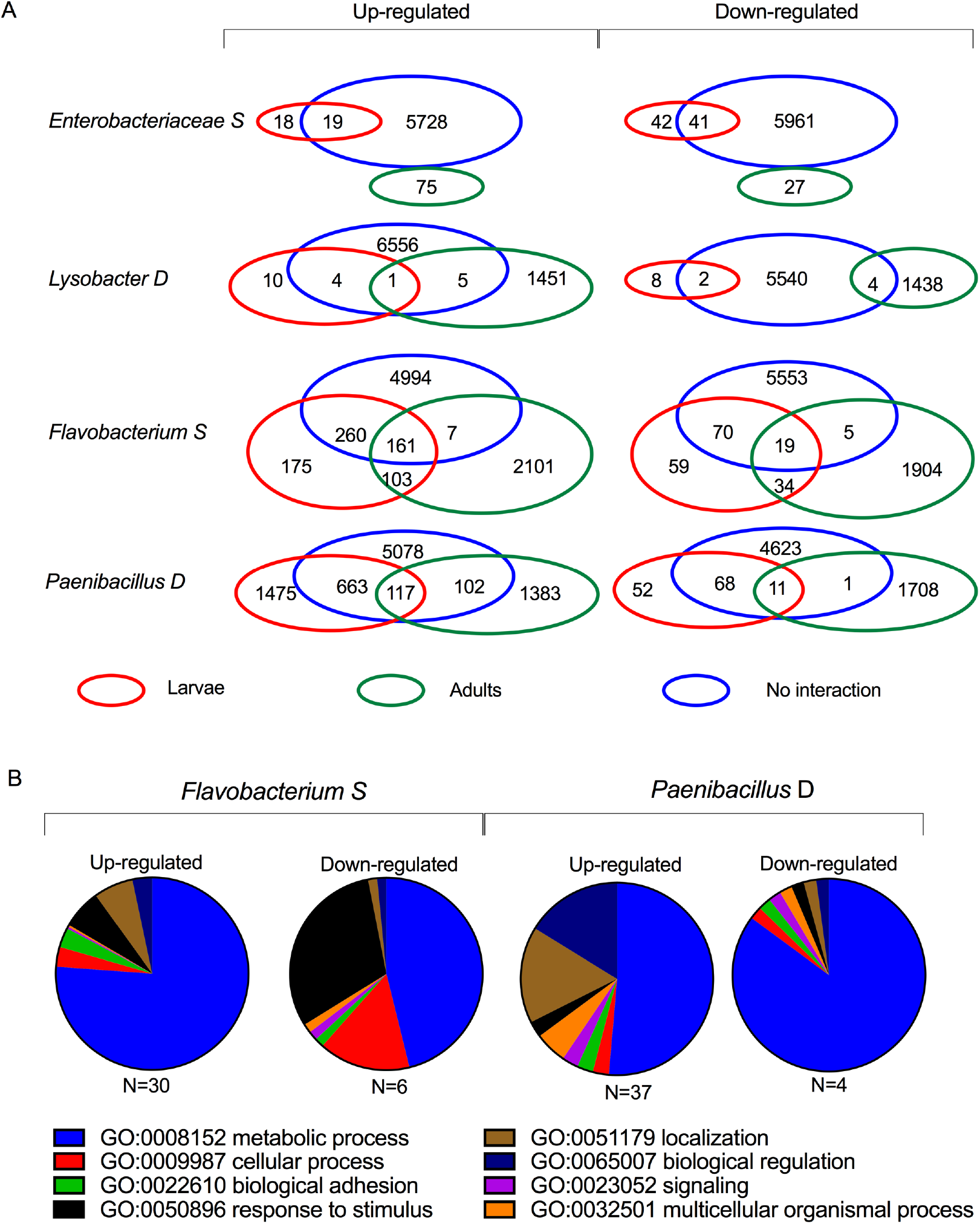
Carry-over effects occur at the level of gene expression for two bacterial isolates. **A**. For each gnotobiotic treatment, the Venn diagrams show the overlap of transcripts that are up- or down-regulated in larvae (green), in adults (red), and that are not associated with a statistical interaction between the two stages (blue). The interaction analysis is provided in Fig. S3. The triple intersection of the Venn diagrams represents the ‘carry-over transcripts’ that were differentially expressed (relative to the non-axenic controls) in the same direction and with a similar magnitude in both larvae and adults from the same gnotobiotic treatment. **B**. The pie charts show the GO categories of ‘carry-over transcripts’ in the *Flavobacterium* S and *Paenibacillus* D gnotobiotic treatments. N indicates the total number of transcripts assigned to functional categories.

### Larval exposure to bacteria results has physiological consequences at the adult stage

To investigate the functional consequences of transcriptional changes associated with gnotobiotic rearing, we examined the metabolic pathways that were differentially regulated at each life stage. We found that almost all the transcripts involved in metabolic pathways that were up-regulated in larvae were related to lipid metabolism (Fig. 5A; Fig. S4). It was also the case in adults for the two bacterial isolates for which we detected a large number of ‘carry-over transcripts’ (*Flavobacterium* S and *Paenibacillus* D). In the *Flavobacterium* S treatment, for instance, we found that most enzymes in the glycerophospholipid metabolism pathways were differentially expressed in larvae (Fig. 5B). To determine whether such transcriptional changes could have functional consequences at the adult stage, we measured the gene expression level of seven triglyceride enzymes in individual adult *Ae. aegypti* females. We found that five and four of these enzymes were indeed up-regulated in the *Flavobacterium* S and *Paenibacillus* D treatments, respectively, relative to the non-axenic controls (Fig. 6A). Moreover, we measured elevated levels of triacylglycerol in adult females from the *Flavobacterium* S and *Paenibacillus* D treatments (Fig. 6B), which translated into higher starvation resistance in these two treatments (Fig. 6C). Thus, these experiments showed that the transcriptional changes detected between mosquitoes exposed to different bacteria at the larval stage were reflected at the functional level in adults.

**Fig. 5.**
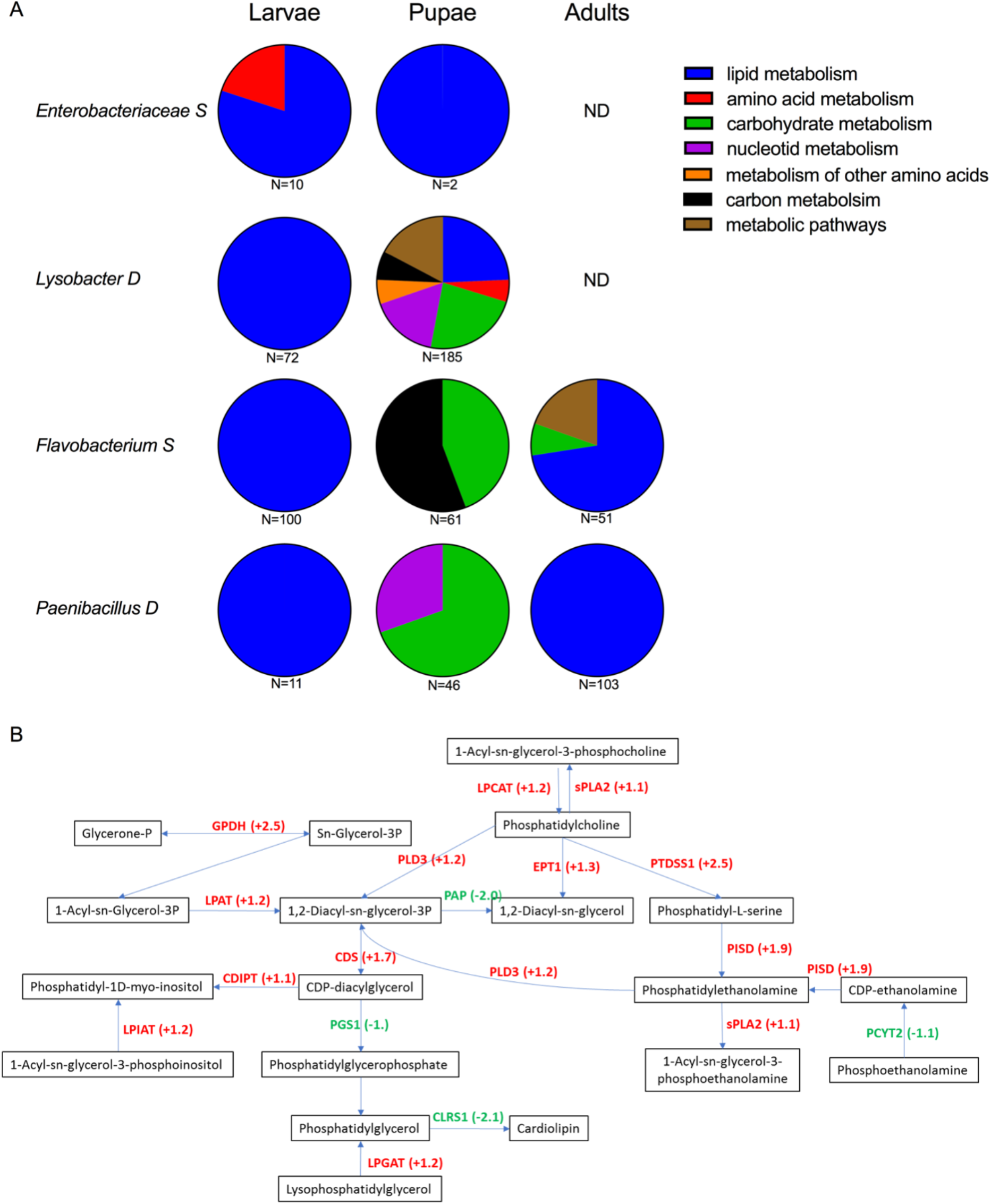
Gene expression changes in gnotobiotic larvae reflect remodeling of lipid metabolism. **A**. The pie charts show the proportion of transcripts involved in the metabolism of different substrates that were up-regulated (relative to the non-axenic controls) in each of the gnotobiotic treatments (in rows) at each life stage (in columns). N indicates the total number of transcripts assigned to each metabolic substrate (ND=not detected). **B**. The flowchart shows significantly up-regulated (red) and down-regulated (green) enzymes in the glycerophospholipid metabolism pathways after *Flavobacterium* S exposure at the larval stage (relative to non-axenic controls). Values in brackets are the transcript log_2_-transformed fold-change of each enzyme in L_4_ larvae. CDIPT: CDP-diacylglycerol--inositol 3-phosphatidyltransferase; LPIAT: lysophospholipid acyltransferase 7; LPGAT: lysophosphatidylglycerol acyltransferase 1; CDS: phosphatidate cytidylyltransferase; GPDH: glycerol-3-phosphate dehydrogenase; LPAT: lysophosphatidate acyltransferase; PLD3: phospholipase D3; PTDSS1: phosphatidylserine synthase 1; PISD: phosphatidylserine decarboxylase; LPCAT: lysophospholipid acyltransferase 5; sPLA2: secretory phospholipase A2; EPT1: ethanolamine phosphotransferase; GPAT3: glycerol-3-phosphate O-acyltransferase 3; CRLS1: cardiolipin synthase ; PGS1: CDP-diacylglycerol--glycerol-3-phosphate 3-phosphatidyltransferase; LPIN1: phosphatidate phosphatase LPIN; PAP: diacylglycerol diphosphate phosphatase / phosphatidate phosphatase; ETNPPL: ethanolamine-phosphate phospho-lyase; PCYT2: ethanolamine-phosphate cytidylyltransferase.

**Fig. 6.**
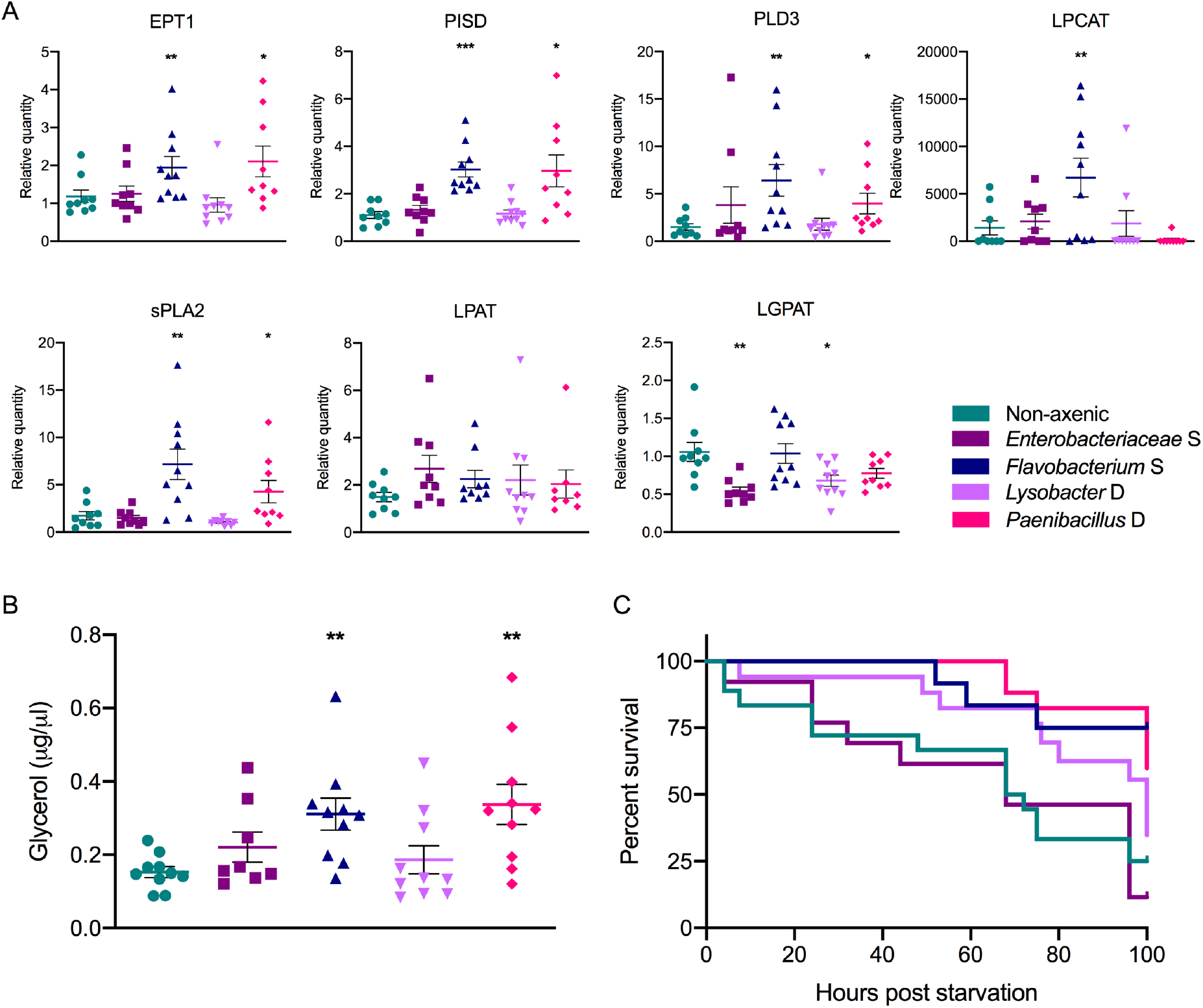
Larval gnotobiotic treatments impact lipid metabolism and resistance to starvation in adult *Ae. aegypti* females. **A**. Bar plots showing the expression levels of triglyceride enzymes in individual adult female mosquitoes (N=9-10 per condition) after larval exposure to *Enterobacteriaceae* S, *Flavobacterium* S, *Lysobacter* D, *Paenibacillus* D or non-axenic control treatment. Expression level was measured by quantitative RT-PCR and normalized to the housekeeping gene *RP49.* EPT1: ethanolamine phosphotransferase; PISD: phosphatidylserine decarboxylase; PLD3: phospholipase D3; LPGAT: lysophosphatidylglycerol acyltransferase 1; sPLA2: secretory phospholipase A2; LPAT: lysophosphatidate acyltransferase and LPCAT: lysophospholipid acyltransferase 5. Asterisks indicate statistically significant differences (\**p*<0.05, \*\**p*<0.005 and \*\*\**p*<0.001) between the gnotobiotic treatment and the non-axenic controls according to a Mann–Whitney *U* test. **B**. Bar plot showing total triacylglycerol (TAG) levels in adult female mosquitoes (N=8-10 pools of 3 individuals per condition) after larval exposure to *Enterobacteriaceae* S, *Flavobacterium* S, *Lysobacter* D, *Paenibacillus* D or non-axenic control treatment. TAG levels were assessed enzymatically and normalized to weight. Asterisks indicate statistically significant differences (\**p*<0.05) between the gnotobiotic treatment and the non-axenic controls according to a Mann– Whitney *U* test. **C**. Kaplan-Meier plots showing adult female survival over time under starvation. Resistance to starvation was measured by monitoring mosquito mortality in triplicate boxes of 10 females at 4- to 8-hour intervals, in two separate experiments. The test statistics of panel **C** are provided in Table S2.

## Discussion

This study builds on our previous work [35] to show that carry-over effects detected at the phenotypic level are associated with massive changes at the molecular level in gnotobiotic *Ae. aegypti* mosquitoes. Specifically, we found that larval exposure to different bacteria results in transcriptomic and metabolic changes that have consequences on adult physiology and fitness. For two of the four gnotobiotic treatments that we examined in more detail (larval exposure to *Flavobacterium* S and *Paenibacillus* D), we noticed that several transcripts related to lipid metabolism were up-regulated in both larvae and adults. We found elevated lipid content in the adult females emerging from these gnotobiotic treatments, which translated into higher starvation resistance. Together, our results provide evidence that carry-over effects in gnotobiotic *Ae. aegypti* reflect trans-stadial metabolic remodeling. This is an important leap forward to understanding the mechanisms driving the effects of mosquito-bacteria interactions at the larval stage on adult fitness.

We first confirmed and expanded the results of our previous study [35] by detecting significant carry-over effects in adult *Ae. aegypti* females after gnotobiotic rearing. We found that exposure to different bacteria at the larval stage influences several traits underlying vectorial capacity, including pupation rate, adult survival, body size and vector competence for DENV. We previously detected such carry-over effects for pupation rate, adult body size and DENV vector competence but not adult survival [35]. This discrepancy could reflect the more rigorous control of inoculum size that we implemented in the present study by adjusting the bacterial concentration to 5 × 10^5^ CFU/ml instead of using a colony of similar size. It could also be due to the larger number of bacterial isolates that were tested (16 vs. three in the previous study) since only seven out of 16 bacterial isolates resulted in a consistent effect on adult survival. The influence of larval exposure to bacteria on adult female survival is epidemiologically meaningful because it is one of the most critical determinants of vectorial capacity [39].

We used an untargeted approach of transcriptome sequencing to examine the gene expression profiles of gnotobiotic larvae, pupae and freshly emerged adults. We detected substantial variation in transcript abundance between the gnotobiotic treatments both in pairwise comparisons and using non-axenic controls as the reference. In addition to the expected differences between life stages, we found that most of the transcriptional changes were related to metabolic processes. This is consistent with other studies suggesting that bacteria play a nutritional role during larval development in mosquitoes [28, 32] and fruit flies [33, 34]. This is also in line with the results from a transcriptional comparison of axenic and gnotobiotic *Ae. aegypti* larvae [27], which found that the lack of mosquito-bacteria interactions at the larval stage results in defects in acquisition and assimilation of nutrients. In contrast, a recent study found that axenic and conventionally reared *Ae. aegypti* only displayed minimal differences in gene expression at the adult stage [41]. Therefore, exposure to different bacteria during larval development could result in more variation in gene expression patterns at the adult stage than the presence of bacteria itself.

A key result of the present study was to identify ‘carry-over transcripts’, that is, transcripts that are differentially regulated in the same direction and with the same magnitude in both larvae and adults. The effect of the larval environment on mosquito adult traits is well established [11–15], but to our knowledge, this is the first report of transcriptional regulation carrying over from larvae to adults. Larval exposure to an *Enterobacter* isolate was shown to modulate immune gene expression in *Ae. aegypti* adults [38], but this did not necessarily imply correlated changes in larval gene expression. Indeed, larval condition can influence the expression of adult immune genes in the absence of changes in larval immunity [42]. We only detected ‘carry-over transcripts’ in two of the four gnotobiotic treatments we focused on (larval exposure to *Flavobacterium* S and *Paenibacillus* D), indicating that carry-over effects observed at the phenotypic level do not necessarily reflect correlated transcript regulation. The molecular mechanisms allowing the trans-stadial persistence of differential gene expression also remain to be elucidated.

Importantly, our study makes a functional link between mosquito-bacteria interactions during larval development, metabolic pathways and the fitness of adults. We found that larval exposure to *Flavobacterium* S and *Paenibacillus* D was associated with remodeling of lipid metabolism, higher levels of lipid storage, higher starvation resistance, and lower DENV titers in adult female *Ae. aegypti*. Flaviviruses are intimately dependent on host lipids and typically reconfigure lipid metabolism in the host cell to create a favorable environment for viral multiplication [43]. The link between lipid metabolism and DENV infection in *Ae. aegypti* was highlighted in several recent studies [44–46] and supports the hypothesis that the observed carry-over effects on DENV vector competence also derive from remodeling of lipid metabolism.

Although gnotobiotic rearing of *Ae. aegypti* larvae with a single bacterial isolate is an over simplification of the biotic environment of wild mosquito larvae, this study provides the proof of principle that the diversity of bacterial communities found in larval breeding sites shapes the physiology and fitness of adult mosquitoes via metabolic changes. In fact, we expect that larval exposure to more complex bacterial communities in natural habitats may drive more pronounced carry-over effects than those observed with a single bacterial isolate. Bacterial taxa found in natural *Ae. aegypti* breeding sites are highly diverse and the majority are not shared across microhabitats [25, 35–37]. This implies that the variable biotic conditions experienced by immature stages likely contribute to create substantial phenotypic variation in adult traits. Because some of these adult traits determine fitness, this phenotypic variation is predicted to translate into significant fitness differences. In turn, these fitness differences are expected to drive the adaptive evolution of female preference for breeding sites that produce the fittest individuals. Surprisingly, few bacterial taxa have been consistently associated with the presence of *Ae. aegypti* larvae in natural breeding sites [36, 47], probably reflecting the functional redundancy between bacterial communities. Thus, advancing our understanding of the molecular and physiological changes that are associated with exposure to different bacterial communities and other microorganisms will improve our understanding of how mosquito-microbiota interactions at the larval stage contribute to shape mosquito fitness.

## Materials and Methods

### Ethics

This study used human blood samples to prepare mosquito artificial infectious blood meals. For that purpose, healthy blood donor recruitment was organized by the local investigator assessment using medical history, laboratory results and clinical examinations. Biological samples were supplied through the participation of healthy adult volunteers at the ICAReB biobanking platform (BB-0033-00062/ICAReB platform/Institut Pasteur, Paris/BBMRI AO203/[BIORESOURCE]) of the Institut Pasteur in the CoSImmGen and Diagmicoll protocols, which had been approved by the French Ethical Committee Ile-de-France I. The Diagmicoll protocol was declared to the French Research Ministry under reference 343 DC 2008-68 COL 1. All adult subjects provided written informed consent.

### Bacterial isolates

All the bacterial isolates used in this study were previously derived from the water of natural *Ae. aegypti* breeding sites in both sylvatic and domestic habitats in Lopé, Gabon in 2014 [35]. A subset of 16 bacterial isolates was initially selected to represent the taxonomical diversity found in both habitats with a balanced proportion of Gram-negative and Gram-positive cultivable bacteria. Based on their *16S* ribosomal RNA gene sequence, the eight selected bacterial isolates from sylvatic breeding sites were taxonomically identified as *Bosea, Cellulomonas, Enterobacteriaceae, Flavobacterium, Leifsonia, Paenibacillus, Pseudoxanthomonas* and *Rahnella.* The eight selected bacterial isolates from domestic breeding sites were identified as *Ancylobacter, Cloacibacterium, Flavobacterium, Leifsonia, Lysobacter, Paenibacillus, Rummeliibacillus* and *Stenotrophomonas*. These bacterial isolates were used to generate the gnotobiotic mosquitoes.

### Gnotobiotic mosquitoes

All mosquitoes used in this study were from the 16^th^ and 17^th^ laboratory generations of an *Ae. aegypti* colony derived from a natural population originally sampled in Thep Na Korn, Kamphaeng Phet Province, Thailand, in 2013. The colony was maintained under controlled insectary conditions (28 ± 1°C; relative humidity, 70 ± 5%; 12h:12h light:dark cycle). Gnotobiotic mosquitoes were generated as previously described [35]. Briefly, eggs were scraped off the blotting paper on which they were laid into a 50-ml conical tube. Eggs were surface sterilized inside a microbiological safety cabinet by sequential incubations in 70% ethanol for 5 min, 3% bleach for 3 min, and 70% ethanol for 5 min, followed by three washes in sterile water. Non-axenic controls were generated by omitting the surface sterilization step. Eggs were allowed to hatch for 1 hour in sterile water in a vacuum chamber, and transferred to sterile 25-ml tissue-culture flasks with filter-top lids and maintained in 15 ml of sterile water. Axenic and non-axenic larvae were seeded to a density of 16 ± 4 per flask. The larvae were fed 60 μl of sterile (autoclaved) fish food (Tetramin) every other day. Axenic L_1_ larvae were made gnotobiotic by adding 5 × 10^5^ CFU/ml of a single bacterial isolate to the flask. Flasks of axenic larvae were maintained for the duration of the experiment and manipulated in the same way to serve as negative controls. Axenic larvae can develop if they are provided a specific diet [28] or maintained in darkness [31], however they do not develop otherwise [26, 35]. Therefore, presence of larvae beyond the first instar in the axenic treatment is indicative of bacterial contamination. Unlike other studies in which the main comparison is between gnotobiotic and axenic treatments, the present study relied primarily on the comparison between different gnotobiotic treatments. During growth, flasks of axenic, non-axenic and gnotobiotic larvae were kept in a cell-culture incubator at 28 ± 1°C under a 12h:12h light:dark cycle. Upon pupation, gnotobiotic and non-axenic mosquitoes were transferred to non-sterile insectary conditions (28°C ± 1°C, 70 ± 5% relative humidity, 12h:12h light:dark cycle). Adults were kept in 1-pint cardboard cups with permanent access to 10% sucrose solution. Adult *Ae. aegypti* females maintained in these insectary conditions were previously shown to share the same gut bacterial microbiota [48].

### Life-history traits

Pupation rate was assessed by counting the number of new pupae on a daily basis. Adult survival was monitored by counting dead mosquitoes daily for 75 days. The adult body size of females was estimated by measuring their wing length [49]. Wings were removed and taped onto a sheet of paper. After scanning the paper, wing lengths were measured from the tip (excluding the fringe) to the distal end of the allula [50] using the Fiji software [51].

### Bacterial load

To quantify the bacterial load of gnotobiotic and non-axenic mosquitoes, individual L_4_ larvae, pupae and newly emerged adults (N=5-23 per condition) were surface sterilized in 70% ethanol for 10 min and rinsed three times in sterile phosphate-buffered saline (PBS). The samples were homogenized in 180 μl of ATL Buffer (Qiagen) with ~20 1-mm glass beads (BioSpec) in a Precellys 24 grinder (Bertin Technologies) for 30 sec at 6,000 rpm. DNA was extracted with the Blood & Tissue kit (Qiagen) following the manufacturer’s instructions. The amount of bacterial DNA was measured by quantitative PCR using the QuantiTect SYBR Green kit (Qiagen) on a LightCycler 96 real-time thermocycler (Roche) following a published method [52, 53]. The *Ae. aegypti* ribosomal protein-coding gene *RP49* (*AAEL003396*) was used for normalization [54]. The relative DNA quantity was calculated as E^-(Cq*_RP49_*-Cq*_16S_*)^, with E being the PCR efficiency of each primer pair. The sequences of all primers used in this study are provided in Table S3.

### Vector competence

Vector competence assays were conducted as previously described [55] using DENV type 1 (DENV-1) isolate KDH0026A [56]. Briefly, 5- to 7-day-old females were deprived of sucrose solution for 24 hours and transferred to a biosafety level-3 facility. They were offered an artificial infectious blood meal for 15 minutes using an artificial membrane-feeding system (Hemotek) with pig intestine as the membrane. The infectious blood meal consisted of a 2:1 mixture of washed human erythrocytes and virus suspension at a final concentration of 7.5 × 10^5^ and 6.0 × 10^5^ focus-forming units (FFU)/ml in two separate experiments, respectively. The blood meal was supplemented with 10 mM adenosine triphosphate to stimulate blood uptake by mosquitoes. Fully engorged females were sorted into 1-pint carton boxes, and kept under controlled conditions (28°C ± 1°C, 70 ± 5% relative humidity, 12h:12h light:dark cycle) in a climatic chamber with permanent access to a 10% sucrose solution. After 14 days of incubation, the head and body of DENV-exposed mosquitoes were separated from each other to determine infection rate (the proportion of blood-fed mosquitoes with a DENV-positive body), dissemination rate (the proportion of infected mosquitoes with a DENV-positive head) and dissemination titer (the amount of infectious virus in the head tissues of DENV-infected mosquitoes). Bodies were homogenized individually in 400 μl of RAV1 RNA extraction buffer (Macherey-Nagel) during two rounds of 30 sec at 5,000 rpm in a TissueLyser II grinder (Qiagen). Total RNA was extracted using the NucleoSpin 96 kit (Macherey-Nagel) following the manufacturer’s instructions. Total RNA was converted to complementary DNA (cDNA) using M-MLV reverse transcriptase (Invitrogen) and random hexamers. The cDNAs were amplified by PCR as described below, using a primer pair targeting the DENV-1 *NS5* gene (Table S3). The heads from DENV-infected bodies were titrated by focus-forming assay in C6/36 cells as previously described [55]. Briefly, heads were homogenized individually in 300 μl of Leibovitz’s L-15 medium supplemented with 2× Antibiotic-Antimycotic (Life Technologies). C6/36 cells were seeded into 96-well plates and incubated 24h to reach sub-confluence. Each well was inoculated with 40 μl of head homogenate in 10-fold dilutions and incubated for 1 hour at 28°C. Cells were overlaid with a 1:1 mix of carboxymethyl cellulose and Leibovitz’s L-15 medium supplemented with 0.1% penicillin (10,000 U/ml)/streptomycin (10,000 μg/ml), 1× non-essential amino acids, 2× Antibiotic-Antimycotic (Life Technologies), and 10% fetal bovine serum (FBS). After three days of incubation at 28°C, cells were fixed with 3.7% formaldehyde, washed three times in PBS, and incubated with 0.5% Triton X-100 in PBS. Cells were incubated with a mouse anti-DENV complex monoclonal antibody (MAB8705, Merck Millipore), washed three times with PBS, and incubated with an Alexa Fluor 488–conjugated goat anti-mouse antibody (Life Technologies). FFU were counted under a fluorescence microscope.

### Transcriptome sequencing

Triplicate pools of 12 L_1_ axenic larvae, 12 L_4_ gnotobiotic larvae, 12 pupae and 12 freshly emerged adult mosquitoes (6 males + 6 females) were surface sterilized with 70% ethanol and washed three times in PBS. They were transferred to a tube containing 800 μl of Trizol (Life Technologies) and ~20 1-mm glass beads (BioSpec). Samples were homogenized for 30 sec at 6,000 rpm in a Precellys 24 grinder (Bertin Technologies). RNA was extracted and purified as previously described [44]. Total RNA was resuspended into 20 μl of RNase-free water, the quality and the quantity of RNA were checked by Nanodrop and Bioanalyzer (Agilent) and stored at –80°C until further use. Sequencing libraries were prepared from pools of 12 individuals using the TruSeq Stranded mRNA library preparation kit (Illumina) following the manufacturer’s instructions. Library quality was checked on a DNA1000 Bioanalyzer chip (Agilent) and quantification made by QuBit DNA HS kit (ThermoFisher). Single-end reads of 65 nucleotides (nt) in length were generated on a HiSeq2500 sequencing platform (Illumina). Reads were cleaned of adapter sequences, and low-quality sequences were removed using Cutadapt version 1.11 [57]. Only sequences ≥25 nt in length were considered for further analysis. STAR version 2.5.0a [58], with default parameters, was used for alignment to the *Ae. aegypti* reference genome (AaegL5.2, https://vectorbase.org). Genes were counted using featureCounts version 1.4.6-p3 [59] in the Subreads package (parameters: −t exon, −g gene_id and −s 1). The number of uniquely mapped reads ranged from 11.3 to 65.6 millions per sample. The raw sequences were deposited to the NCBI Gene Expression Omnibus repository under accession number GSE173472.

### Differential gene expression

All analyses of transcript abundance were performed using R version 3.6.1 [60] and the Bioconductor package DESeq2 version 1.24.0 [61]. Normalization and dispersion estimation were performed with DESeq2 using the default parameters and statistical tests for differential expression were performed applying the independent filtering algorithm. Differential expression between treatments for larval, pupal and adult stages was tested with a generalized linear model. For each pairwise comparison, raw *p* values were adjusted for multiple testing according to the Benjamini and Hochberg procedure [62]. Genes with both an absolute log_2_-transformed fold-change (log2FC) >1 and an adjusted *p* value <0.05 were considered significantly differentially expressed. Both up- and down-regulated transcripts (relative to the non-axenic controls) were imported into the gene ontology (GO) enrichment analysis tool based on protein analysis through the evolutionary relationships classification system. The molecular functions, cellular component, and biological process GO categories were considered significantly enriched and represented in the pie charts when their adjusted *p* values were <0.05. Up-regulated transcripts were also imported into the pathway enrichment analysis tool of the Kyoto Encyclopedia of Genes and Genomes (KEGG) pathway database.

### Lipid content

Lipid content was evaluated in 3-day-old adult *Ae. aegypti* females following a published method [63]. Females (N=8-10) were individually homogenized in 125 μl of TET buffer (10 mM Tris pH 8, 1 mM EDTA, 0.1% Triton X-100) and ~20 1-mm glass beads (BioSpec) in a Precellys 24 grinder (Bertin Technologies) for 30 sec at 6,000 rpm. The samples were centrifuged for 3 min at 14,000 rpm and 50 μl of the supernatant was incubated at 72°C for 30 min. The triacylglyceride content was assessed enzymatically using the Free Glycerol Reagent (F6428 kit, Sigma) and quantified using a glycerol standard (G7793, Sigma), following manufacturer’s instructions, and normalized to mosquito fresh weight.

### Starvation resistance

Starvation resistance was evaluated in 3- to 5-day-old adult females following a published method [64]. Triplicate groups of 10 females from each condition (gnotobiotic and non-axenic treatments) were maintained in pint-cardboard cups. The sucrose solution was removed and the number of dead mosquitoes was monitored at 4- to 8-hour intervals for 100 hours.

### Statistics

Statistical analyses were performed with JMP version 14.0 (www.jmpdiscovery.com) and GraphPad Prism version 9.0 (www.graphpad.com). To analyze pupation rate, the 50% pupation day (PD_50_) was estimated using the Kaplan-Meier estimator for each replicate flask of each experiment. PD_50_ values were compared between treatments by analysis of variance (ANOVA) weighted by the sample size. To account for uncontrolled variation between experiments, wing lengths were normalized to the non-axenic controls of the same experiment and compared between treatments by ANOVA. Survival curves were analyzed as a function of the treatment, the experiment (if >1) and their interaction using a Cox model and a Wald test. The interaction was removed from the final model when non-significant. Bacterial loads and individual gene expression levels were compared with Mann–Whitney *U* test. DENV infection and dissemination rates were analyzed by logistic regression as a function of the treatment, the experiment and their interaction, followed by likelihood-ratio χ^2^ tests. Dissemination titer was log_10_-transformed and analyzed by ANOVA as a function of the treatment, the experiment and their interaction.

## Acknowledgments

We thank Catherine Lallemand for the maintenance of mosquito colonies and the Lambrechts lab members for their insights. We are grateful to Alongkot Ponlawat for the initial collection of *Ae. aegypti* in Thailand. We thank Christophe Paupy, Davy Jiolle, Diego Ayala, Amine Ghozlane, Marc Monot, Karima Zouache, Edwige Martin, Vincent Raquin and Guillaume Minard for their input and assistance at earlier stages of the study. This work was primarily funded by the Agence Nationale de la Recherche (grant ANR-16-CE35-0004-01). It was also supported by the French Government’s Investissement d’Avenir program Laboratoire d’Excellence Integrative Biology of Emerging Infectious Diseases (grant ANR-10-LABX-62-IBEID to LL), and the City of Paris Emergence(s) program in Biomedical Research (to LL). The Biomics platform was supported by France Génomique (ANR-10-INBS-09-09) and IBISA. The funders had no role in study design, data collection and interpretation, or the decision to submit the work for publication.

## Author contributions

EG, LBD, CVM and LL conceived of the study. LBD collected and isolated the bacteria. EG, FA and SD carried out the mosquito experiments. OS prepared and sequenced the libraries. RL processed the sequencing reads. HV performed the differential gene expression analysis. EG and LL analyzed the experimental data and wrote the manuscript. All authors approved the final version of the manuscript.

## Supporting Information

**Suppl. Table 1.**
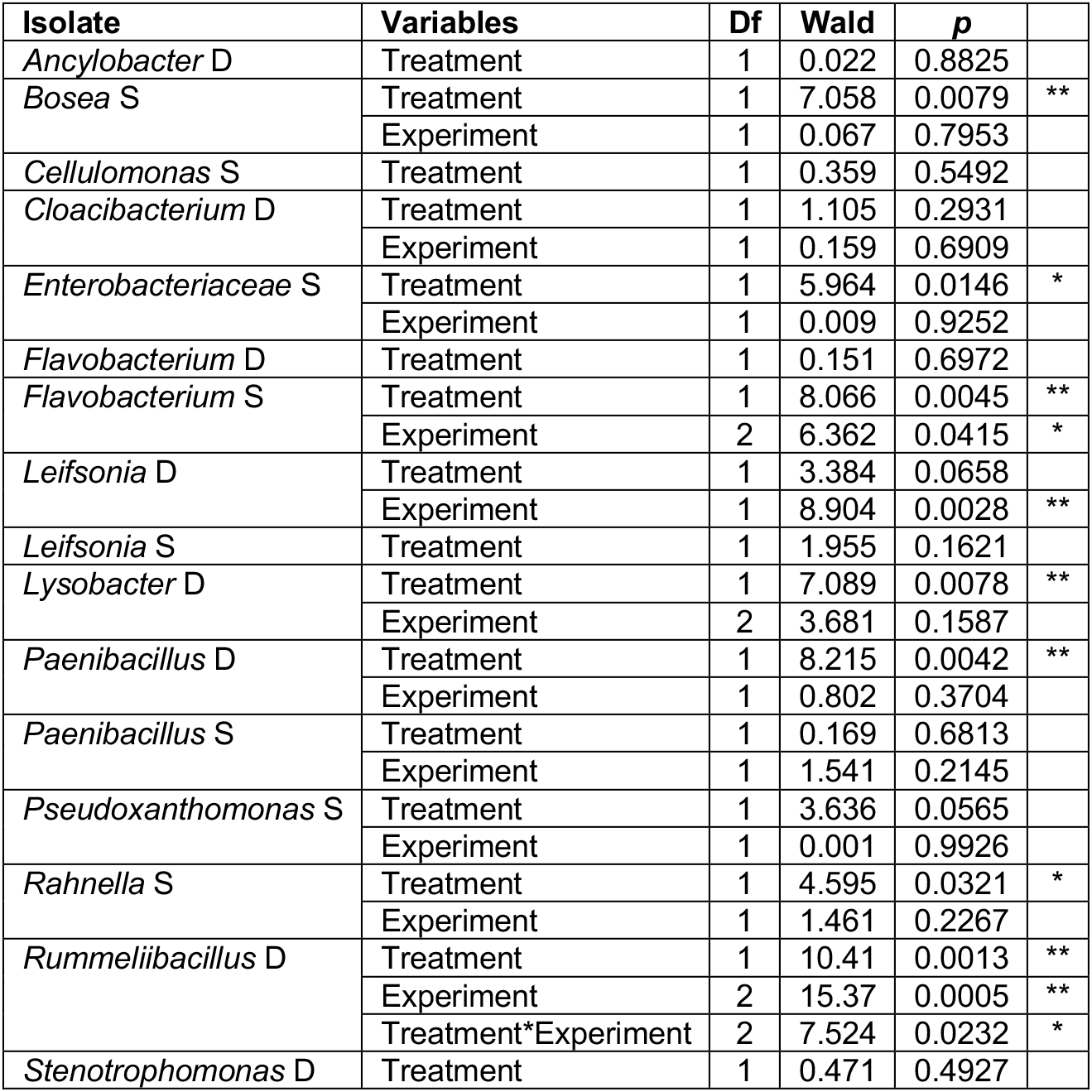
Test statistics of adult female survival analysis after larval exposure to different bacterial isolates. The table shows the statistical analyses of survival curves displayed in Fig. 1C. Survival data were analyzed by Wald test based on the Cox model. For each isolate, the initial model included the treatment (gnotobiotic vs. non-axenic), the experiment (if more than one) and their interaction. The interaction term was removed from the model if non-significant. Df: degrees of freedom. Asterisks indicate statistically significant differences (\**p*<0.05, \*\**p*<0.005).

**Suppl. Table 2.** Test statistics of adult female survival analysis under starvation. The table shows the statistical analysis of survival curves displayed in Fig. 6C. Survival data were analyzed by Wald test based on the Cox model. For each isolate, the initial model included the treatment (gnotobiotic vs. non-axenic), the experiment (up to three) and their interaction. The interaction term was removed from the model if non-significant. Df: degrees of freedom. Asterisks indicate statistically significant differences (\**p*<0.05, \*\**p*<0.005).

| Isolate | Variables | Df | Wald | $p$ | |
| --- | --- | --- | --- | --- | --- |
| *Enterobacteriaceae* S | Treatment | 1 | 0.538 | 0.4634 |  |
|  | Experiment | 1 | 2.284 | 0.1307 |  |
| *Flavobacterium* S | Treatment | 1 | 4.727 | 0.0297 | * |
|  | Experiment | 1 | 9.079 | 0.0026 | ** |
|  | Treatment*Experiment | 1 | 3.911 | 0.0480 | * |
| *Paenibacillus* D | Treatment | 1 | 6.953 | 0.0084 | ** |
|  | Experiment | 1 | 9.625 | 0.0019 | ** |
|  | Treatment*Experiment | 1 | 3.933 | 0.0473 | * |
| *Lysobacter* D | Treatment | 1 | 2.493 | 0.1144 |  |
|  | Experiment | 1 | 5.972 | 0.0145 | * |

**Suppl. Table 3.**
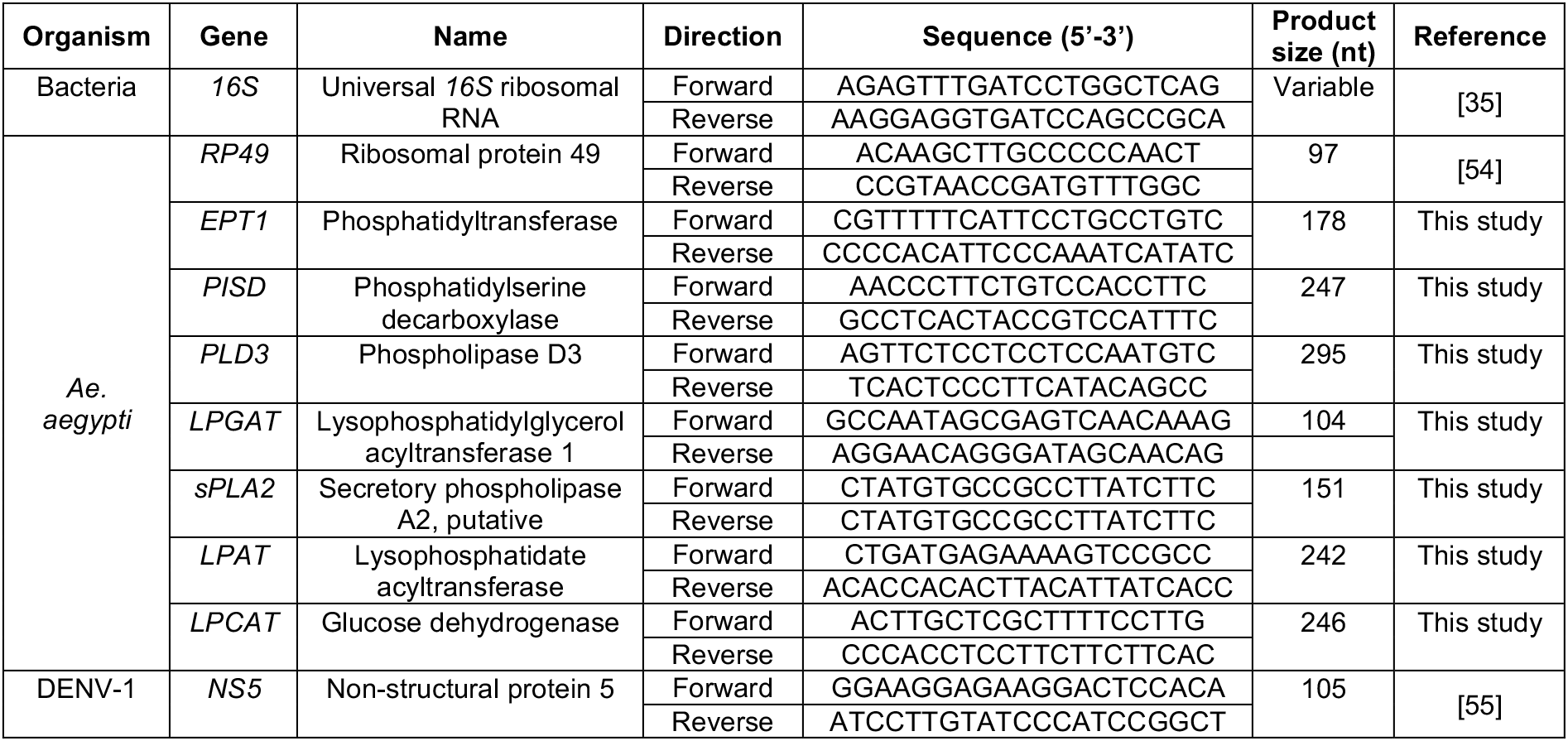
List of primers used in this study.

**Suppl. Fig. 1.**
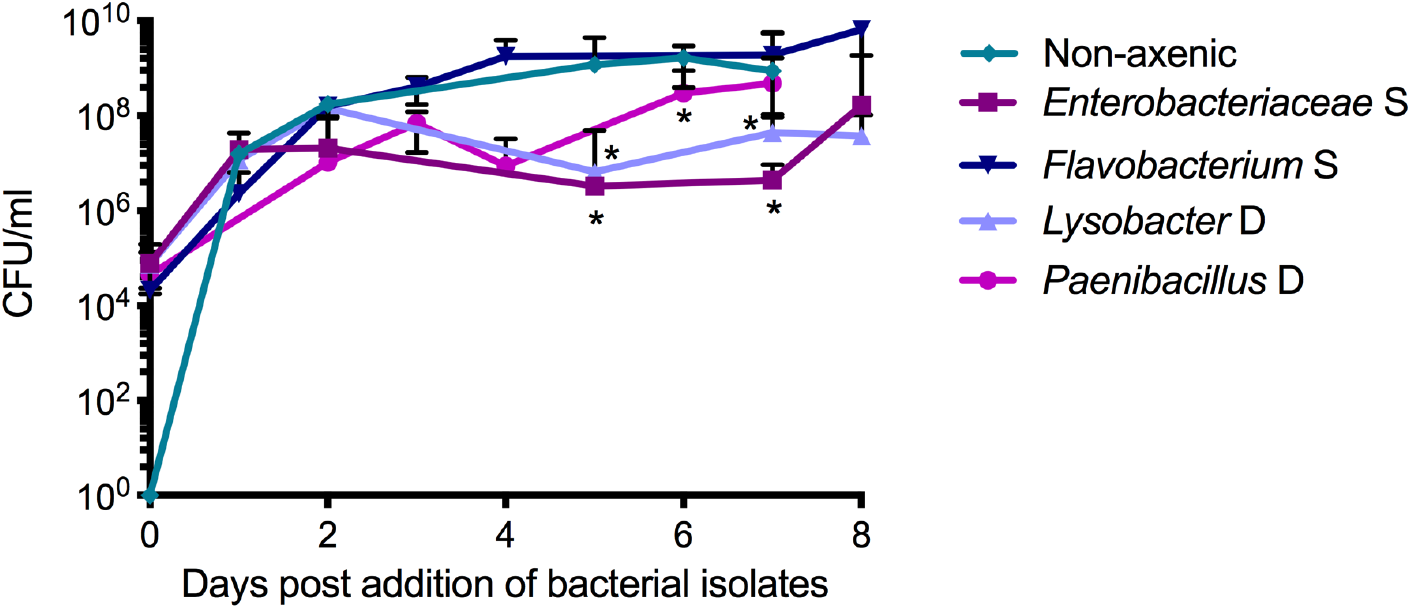
Time course of bacterial concentration in the larval rearing water. The concentration of cultivable bacteria in the rearing flasks was estimated by counting the number of bacterial colonies that grew on LB plates from larval rearing water samples for each gnotobiotic treatment and for the non-axenic control treatment. Data are expressed as colony-forming units (CFU)/ml of water (mean ± SEM of triplicates) on a log_10_ scale. Sterile water was added on day 0, as verified by the absence of CFU in the non-axenic control treatment. Asterisks indicate statistically significant differences between the non-axenic and gnotobiotic treatments (\**p*<0.05) by Mann–Whitney *U* test at each time point.

**Suppl. Fig. 2.**
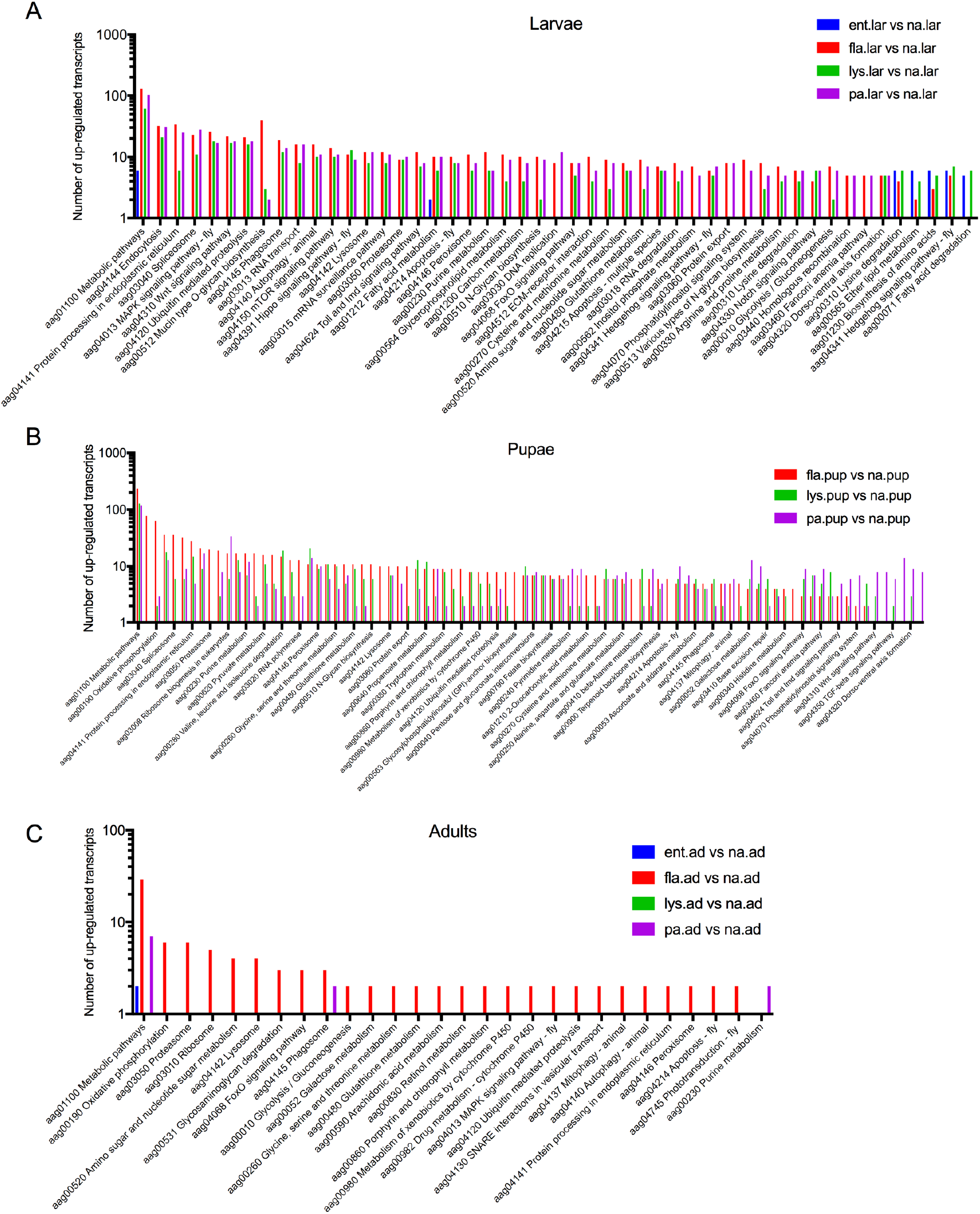
Pathway enrichment analysis of up-regulated transcripts. The number of up-regulated transcripts (relative to the non-axenic controls) in larvae (**A**), pupae (**B**) and newly emerged adults (**C**) is shown for each gnotobiotic treatment (represented by a different color) as a function of Kyoto Encyclopedia of Genes and Genomes (KEGG) categories. In panel **B**, the ent.pup vs. na.pup comparison was not represented by a large enough number of transcripts to perform the pathway enrichment analysis.

**Suppl. Fig. 3.**
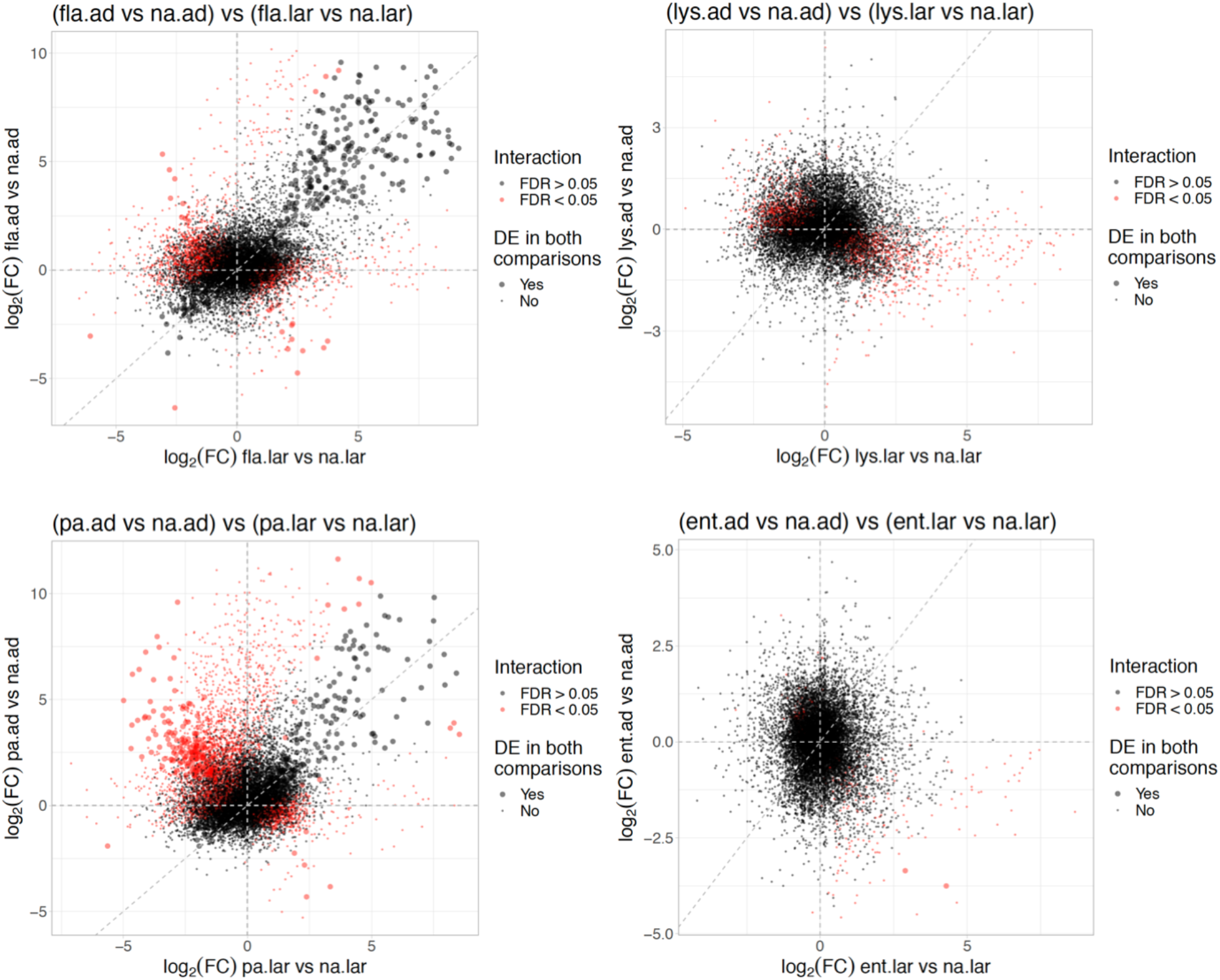
Interaction analysis between larval and adult expression. For all detected transcripts, the log_2_-transformed fold-change (log2FC) in adults (*y*-axis) is plotted as a function of the log2FC in larvae (*x*-axis), stratified by gnotobiotic treatment. The large dots represent transcripts that were differentially expressed (relative to non-axenic controls) both in larvae and in adults. The red dots represent transcripts with a statistically significant interaction (false discovery rate [FDR]<0.05) between adults and larvae whereas the grey dots represent transcripts with a non-significant interaction (FDR>0.05).

**Suppl. Fig. 4.**
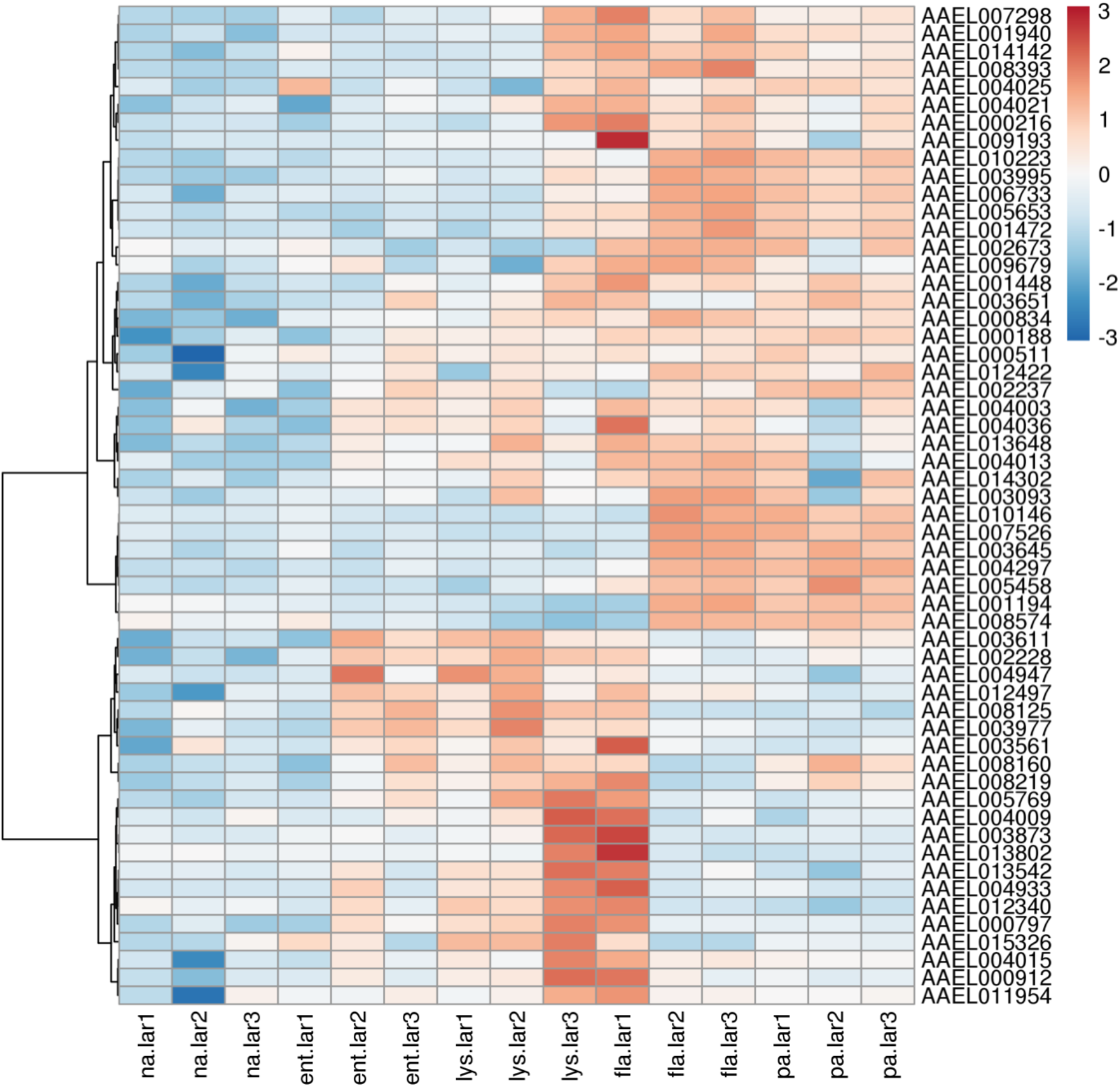
Transcriptional profiles of lipid metabolism genes. The heatmap shows the standardized expression level of differentially expressed transcripts involved in lipid metabolism at the larval stage. Transcripts are identified on the right side by their gene ID. Each column in the heatmap represents a sample, in triplicate for each gnotobiotic treatment. The heatmap is based on the variance-stabilized transformed count matrix. Rows have been re-ordered by hierarchical clustering (shown by the tree on the left) using the correlation distance and the Ward aggregation criterion. The color scale represents the row-centered expression level of each transcript.

